# THE HYBRID HISTORY OF ZEBRAFISH

**DOI:** 10.1101/2024.06.10.598382

**Authors:** Braedan M. McCluskey, Peter Batzel, John H. Postlethwait

**Affiliations:** Minnesota Supercomputing Institute, University of Minnesota, Minneapolis, MN 55455; Institute of Neuroscience, University of Oregon, Eugene, OR 97403-1254

**Keywords:** Zebrafish, Danios, Phylogenomics, Genome Structure, Hybrid Species, Introgression

## Abstract

Since the description of the zebrafish *Danio rerio* in 1822, the identity of its closest living relative has been unclear. To address this problem, we sequenced the exomes of ten species in genus *Danio* and used them to infer relationships across the 25 chromosomes of the zebrafish genome. The majority of relationships within *Danio* were remarkably consistent across all chromosomes. Relationships of chromosome segments, however, depended systematically upon genomic location within zebrafish chromosomes. Regions near chromosome centers identified *D. kyathit* and/or *D. aesculapii* as the closest relative of zebrafish, while segments near chromosome ends supported only *D. aesculapii* as the zebrafish sister species. Genome-wide comparisons of derived character states revealed that danio relationships are inconsistent with a simple bifurcating species history and support an ancient hybrid origin of the *D. rerio* lineage. We also found evidence of more recent gene flow limited to the high recombination ends of chromosomes and several megabases of chromosome 20 with a history distinct from the rest of the genome. The additional insight gained from incorporating genome structure into a phylogenomic study demonstrates the utility of such an approach for future studies in other taxa. The multiple genomic histories of species in the genus *Danio* have important implications for comparative studies in these species and for our understanding of the hybrid evolutionary history of zebrafish.

## Introduction

Despite the status of *Danio rerio* as a major model organism, its recent evolutionary history remains unclear. Different phylogenetic studies of the genus *Danio* pointed to several different species as the sister group to zebrafish (Meyer, et al. 1993; Fang 2003; Quigley, et al. 2004; Mayden, et al. 2007; Mayden, et al. 2008; Fang, et al. 2009; Tang, et al. 2010; McCluskey and Postlethwait 2015) due in part to limited phylogenetic signal in the few genes examined and the ongoing description of danio biodiversity (Fang 1998; Kullander and Fang 2009a). The first study of genus *Danio* using a phylogenomic approach (McCluskey and Postlethwait 2015) proposed that the discordant results of past studies could all be due in part to the multiple genomic histories contained within the genomes of danios.

Determining relationships among closely related species relies on genetic variants in the form of shared derived characters (SDCs) ultimately arising from ancestral alleles (denoted “A”) being converted to derived alleles (denoted “B”) via *de novo* mutation in an ancestral population. A single population history can contain several different genomic histories (Fig. 1A-C). Discordant histories can arise from various processes that can be discerned by the relative frequencies of derived characters shared by different taxa across the genome. When species arise rapidly from an ancestral population, alleles segregating in the ancestral population can be inherited in a pattern that does not necessarily reflect the history of population bifurcation (Holder, et al. 2001) in a process known as incomplete lineage sorting of alleles (ILS, Fig. 1D). Under ILS, the most frequent pattern of SDCs will be those shared by sister species, as new alleles are generated in an ancestral population shared by only two species (dashed lines in Fig. 1D). Alleles inherited by ILS will also occur, but will be balanced between sister taxa, with each sister species sharing roughly the same number of alleles with more distantly related species (Fig. 1D).

**Figure 1:**
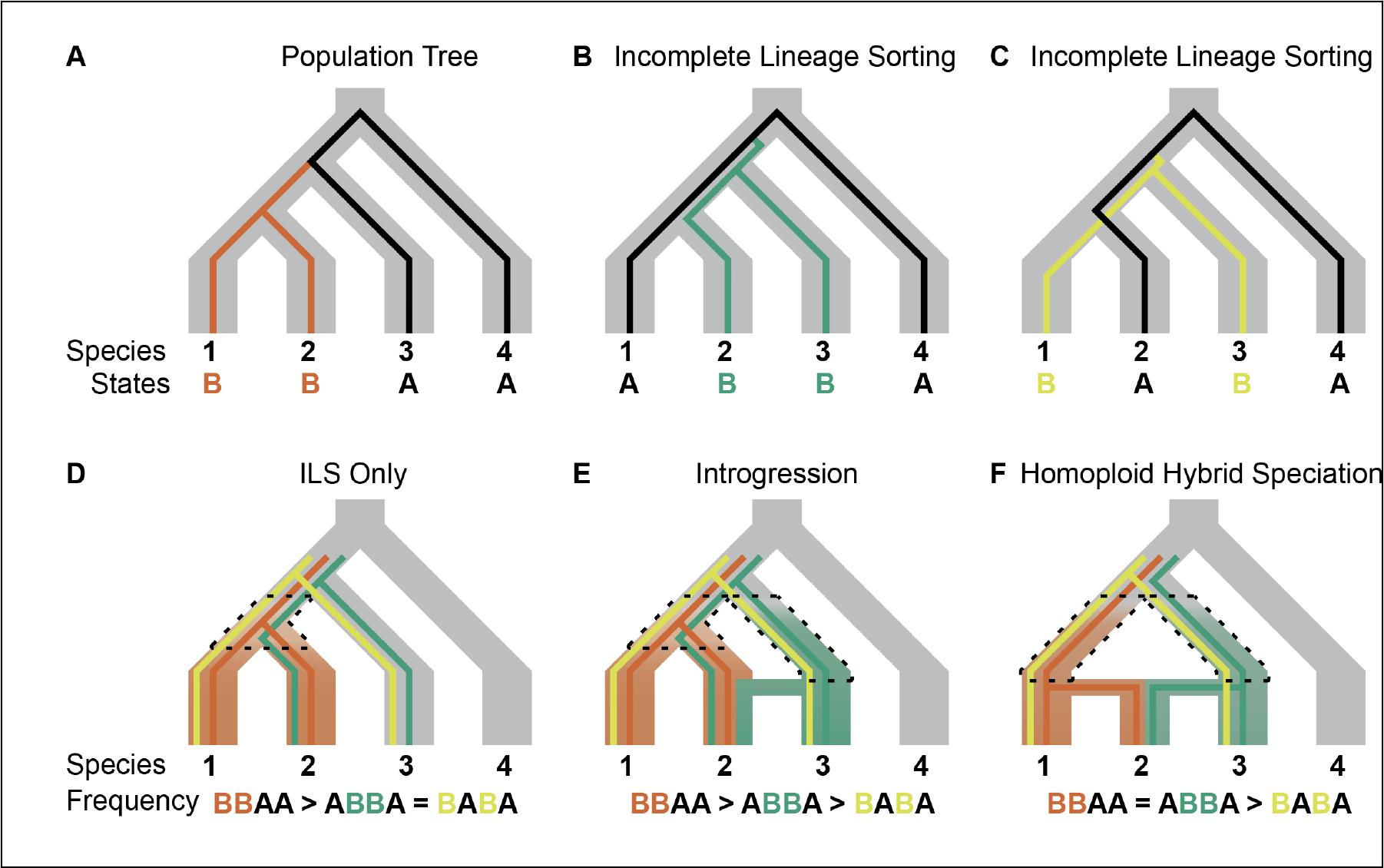
Demographic events and distribution of shared derived characters in related lineages. Ancestral alleles are denoted “A” and derived alleles “B”, with alleles segregating in multiple individuals in populations over time. Derived alleles present in the ancestral population of the three species are shown as colored lines. Alleles arising in ancestral populations after the split from the outgroup and shared by only two species (dashed lines) are shown with shading. (A) Mutations can occur and reflect the history of the population tree. (B, C) Alternative ways of inheriting ancestrally segregating polymorphisms by incomplete lineage sorting of alleles. (D) Expectations with ILS as the only source of multiple genomic histories (compare to panels B and C). (E) Expectations with introgression from one species. Introgression can also occur from two different species, not shown. (F) Expectations with homoploid hybrid speciation. Homoploid hybrid speciation is a special type of hybridizationassociated speciation.

Alleles can also be exchanged between diverged populations via introgression (Fig. 1E). Introgression can be inferred from an excess of one pattern of alleles that can be inherited by ILS. The regions of the genome affected by introgression can be detected by an excess of derived character states relative to other regions in the genome (Durand, et al. 2011; Eaton and Ree 2013; Supple, et al. 2013; Martin, et al. 2015a; Li, et al. 2016). Regions of high recombination are expected to preferentially retain alleles passed by introgression because they can be more easily separated from linked maladaptive alleles than alleles in low-recombination regions. The most extreme example of discordant genomic histories occurs in instances of hybridization-associated speciation (HAS), such as when a new species forms following a hybrid swarm, repeated gene flow following initial divergence, or via homoploid hybrid speciation (HHS) (Fig. 1F). Different forms of HAS can be difficult or impossible to distinguish using only genomic data (Schumer, et al. 2014), but species originating from any type of HAS are more closely related to each parent species than to any other species and should harbor roughly equal SDCs inherited from each parent species. In contrast to introgression, signals of HAS will occur genome wide rather than mainly in regions with high rates of recombination.

Phylogenomic analyses of several taxa have used patterns of SDCs across the genome to infer population histories from local genome histories. Such groups include systems with ecology and mating systems predisposing them to gene flow, such as oaks, broomrape, cats and mosquitoes (Eaton and Ree 2013; Hipp, et al. 2014; Fontaine, et al. 2015; Li, et al. 2016), groups that radiated rapidly such as cichlid fish, swordtails, tomatoes, and birds (Meyer, et al. 2015; Prum, et al. 2015; Pease, et al. 2016; Schumer, et al. 2018), and key model organisms including drosophila (Clark, et al. 2007; Garrigan, et al. 2012) and primates (Scally, et al. 2012; Ting and Sterner 2013; Prufer, et al. 2014; Sankararaman, et al. 2014; Gordon, et al. 2016).

Several aspects of genome structure have been shown to correlate with incomplete lineage sorting and/or introgression including chromosomal location, gene density, and recombination rate (Mallet 2005; Turner, et al. 2005; Ellegren, et al. 2012; Scally, et al. 2012; Burri, et al. 2015; Fontaine, et al. 2015). These effects can result in distinct patterns across the genome, which are not apparent without incorporating genomic structure into the analysis.

As more research groups use the genus *Danio* as a model for evolution (Mahalwar, et al. 2014; McMenamin, et al. 2014; Patterson, et al. 2014; McCluskey and Postlethwait 2015; Singh and Nusslein-Volhard 2015; Irion and Nusslein-Volhard 2019; Podobnik, et al. 2020; Huang, et al. 2021; McCluskey, Liang, et al. 2021; McCluskey, Uji, et al. 2021; Toomey, et al. 2022; Podobnik, et al. 2023), it becomes increasingly important to understand relationships within the genus to properly interpret the trajectory of evolutionary change. Here, we demonstrate how genome structure mediated admixture between the closest relatives of zebrafish. Our findings show that the *Danio rerio* lineage arose from genome-wide admixture between two separate lineages, compatible with homoploid hybrid speciation.

## Results

### Exome enrichment and data validation

To investigate the evolutionary history of the genus *Danio*, we used RNA baits complementary to annotated *D. rerio* nuclear mRNA genes to isolate and sequence exomes from ten danio species varying in body size, pigment pattern, barbel number, and geographic distribution, as well as one species from the closely related *Devario* genus (Supplemental Table S1). We applied stringent filtering to ensure that only high-quality sites were used for phylogenetic inference (see Materials and Methods). Our main data set comprised 15.44 megabases of aligned nucleotides from 11,682 genes (1.1% of the nuclear genome sampled from nearly half of all protein-coding genes with annotated primary transcripts), which is over 100 times more sequence than the most recent *Danio* phylogenomic study based on reduced representation sequencing (McCluskey and Postlethwait 2015). This data set has notable limitations, however. Capture baits lacked mitochondrial genes, so these data can only address the history of the nuclear genome. Furthermore, filtering for codons with alignable sequence across all species means our data set will miss sequences that had identity too low to effectively bind to the RNA baits, a possibility for some genes given the evolutionary distances involved.

**Supplemental Table S1.** Illumina library statistics.

| Species | Total<br>Reads | Reads Aligned to<br>GRCz10 | Proportion<br>Aligned | Exome<br>Coverage |
| --- | --- | --- | --- | --- |
| *Danio rerio* | 41603616 | 41292106 | 0.993 | 87.839 |
| *Danio aesculapii* | 35171505 | 31812385 | 0.904 | 67.673 |
| *Danio kyathit* | 50412319 | 45793130 | 0.908 | 97.414 |
| *Danio nigrofasciatus* | 37389413 | 33110288 | 0.886 | 70.434 |
| *Danio albolineatus* | 31916307 | 27771158 | 0.870 | 59.077 |
| *Danio kerri* | 39820645 | 34829199 | 0.875 | 74.091 |
| *Danio choprae* | 52038977 | 45517922 | 0.875 | 96.829 |
| *Danio erythromicron* | 25080033 | 20885438 | 0.833 | 44.429 |
| *Danio margaritatus* | 35758836 | 30482592 | 0.852 | 64.845 |
| *Danio meghalayensis* | 39328467 | 33221487 | 0.845 | 70.671 |
| *Devario aequipinnatus* | 31491855 | 22986682 | 0.730 | 48.899 |

To validate the accuracy of this exome-sequencing method in recapitulating known sequences for each species and to determine its utility for constructing gene trees, we compared sequences generated by our pipeline to reference sequences for *rag1* and *rho*, two genes used in previous phylogenetic studies with sequences for danios and related species available in GenBank. The reconstituted *rho* and *rag1* sequences from our exome data all had BLAST hits to the correct species (when present in GenBank) with greater than 98% identity (Supplemental Table S2).

**Supplemental Table S2:**
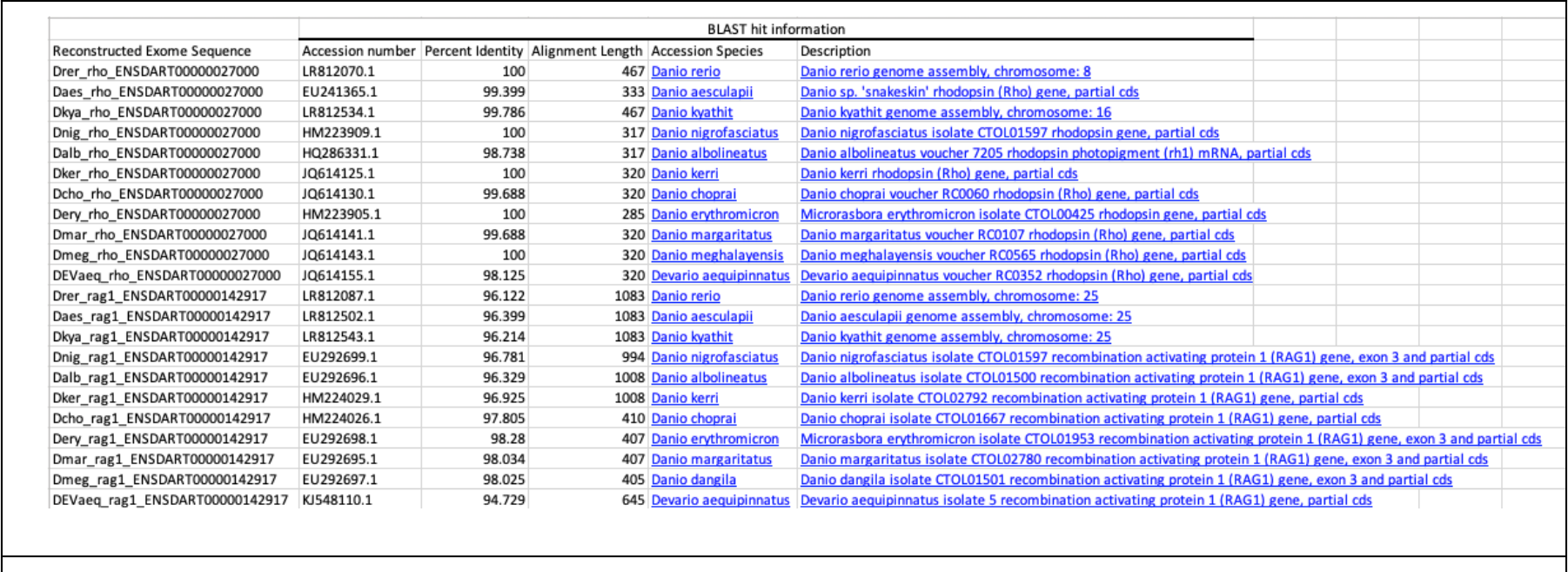
BLAST results for *rho* and *rag1* nucleotide sequences for *Danio* and *Devario* available in public databases, when compared to sequences reconstructed in this study, validate species identification and method robustness.

### Widespread Genealogical Incongruence

To investigate relationships among danio species, we inferred maximum likelihood phylogenies using several partitioning strategies to explicitly account for genome structure, inheritance patterns, and multiple genomic histories. As a first approximation of the history of these species, we assembled all codons that passed filtering into a single alignment and inferred a single phylogeny from genes sampled from the entire nuclear genome (Fig. 2a). We refer to this phylogeny as the ‘unpartitioned genome phylogeny’.

**Figure 2:**
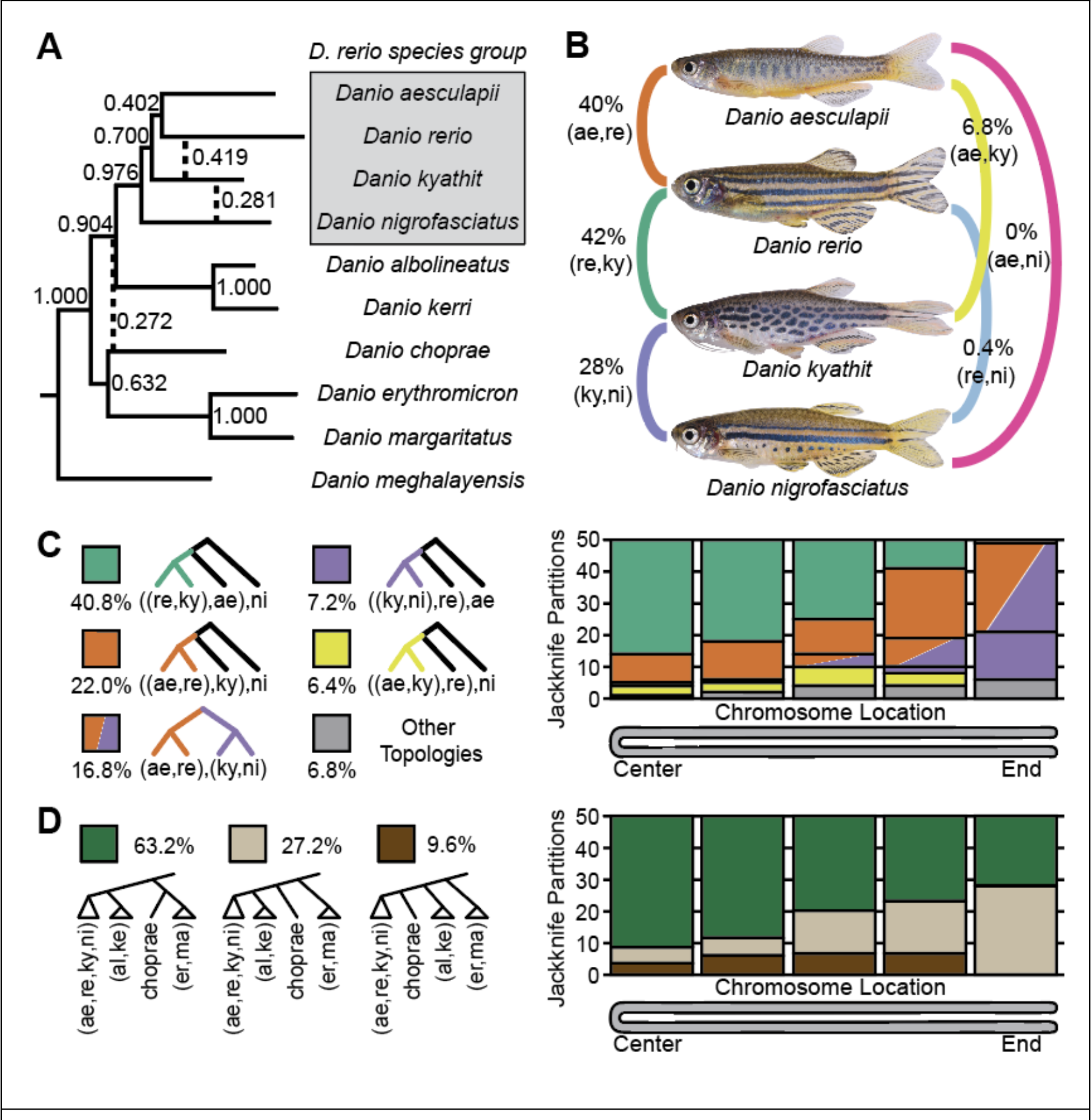
Genealogical discordance and the effects of chromosome structure. (A) The unpartitioned genome phylogeny inferred from the concatenated sequence for each species with concordance factors based on the phylogenies inferred for the 250 jackknife partitions. Dashed lines denote relationships not found in the unpartitioned genome topology but supported by at least 10% of jackknife partitions. Splits supported by <10% of trees are not shown. A gray box demarcates the *D. rerio* species group. (B) Concordance factors of pair-wise splits grouping two species as sister taxa within the *D. rerio* species group for the 250 jackknife partitions, with major concordances shown on the left and minor concordances listed on the right. (C, left) Five genealogical histories in the *D. rerio* species group explain relationships inferred for 93.2% of windows across the genome and a minority of other topologies explain the rest (6.8%, gray). (C, right) The frequency of the most common jackknife topologies binned according to their position across chromosomes, indicated as a folded chromosome with the two telomeres at the right and the chromosome middle on the left. Each bin contains 50 partitions corresponding, for example, to alignments near the ends of each of the 25 *Danio* chromosomes. The inferred sister species of *D. rerio* depends on the position of the partition along the chromosome, with the *D. rerio* – *D. kyathit* relationship (re, ky; green, see panel C, left) supported across the middle of most chromosomes, while the *D. rerio – D. aesculapii* relationship (ae, re; rust) supported near the telomeres of most chromosomes. Note also that few (ky, ni; purple) trees occurred at chromosome centers and many (ky, ni; purple) trees occupied partitions near the telomeres. (D, left) The relationship of *D. choprae* relative to other members of the genus phylogeny varied according to chromosome position. ((D, right) The centers of chromosomes supported the placement of *D. choprae* inferred by the unpartitioned genome phylogeny (green), while chomosome ends showed more variation in the relationships supported.

The phylogeny inferred from the unpartitioned genome-wide nuclear dataset strongly supported the monophyly of the *D. rerio* species group (*sensu* Fang, 1998). This group is represented in this study by *D. rerio, D. aesculapii*, *D. kyathit* and *D. nigrofasciatus*. Other members of this group not sampled in this study likely include *D. quagga* (formerly referred to as *D*. aff. *kyathit*) and *D. tinwini* (McCluskey and Postlethwait 2015). This genome-wide inferred phylogeny placed *D. aesculapii* as the sister species to *D. rerio*, with small internal branches placing *D. kyathit* and *D. nigrofasciatus* as diverging more basally (Fig. 2A). Outside of the *D. rerio* species group, relationships were consistent with previous studies. Two species — *D. albolineatus* and *D. kerri* — fell just outside the *D. rerio* species group, followed by a group of three species — *D. choprae*, *D. erythromicron* and *D. margaritatus* — with the large-bodied *D. meghalayensis* diverging at the base of the genus (Fig. 2A).

To test how much of the danio genome supported the unpartitioned genomic topology, we used a jackknife subsampling approach to divide the nuclear genome into 250 windows (ten windows for each of the 25 chromosomes) based on chromosome position in the *D. rerio* reference genome. Each window had the same number of nucleotides of filtered, aligned sequence as the other nine windows on that chromosome with jackknife windows ranging in size from 37 kb on chromosome 22 to 84 kb on chromosome 7 due to chromosomes varying in sequence length.

The chromosome-level structure of the zebrafish genome is a good approximation of the genomes of other danios. Like most species in family Cyprinidae, zebrafish has 25 pairs of chromosomes ranging from metacentric to subtelocentric (Pijnacker and Ferwerda 1995; Daga, et al. 1996; Amores and Postlethwait 1999; Gornung, et al. 2000; Sola and Gornung 2001; Traut and Winking 2001; Phillips, et al. 2006) with increased recombination rates near their ends (Singer, et al. 2002; Bradley, et al. 2011; Anderson, et al. 2012; Howe, et al. 2013; Wilson, et al. 2014a). Translocations between different chromosomes are rare in cyprinids as demonstrated by comparisons of gene order between zebrafish and distantly related Cyprinids, including carp, goldfish, and grass carp (Xu, et al. 2014; Wang, et al. 2015; Chen, et al. 2019). Intrachromosomal rearrangements, however, have occurred in cyprinids (Avise and Gold 1977), with some even between *D. rerio* strains (Freeman, et al. 2007). Using our jackknife strategy, we could infer the histories of physically linked genomic windows as well as determine the effects of recombination rate throughout the genome.

Nearly all relationships found in the unpartitioned genomic phylogeny were supported by more than 90% of the 250 jackknife windows (Fig. 2A) demonstrating the sufficiency of the jackknife windows to robustly infer phylogenetic relationships. It is striking, therefore, that three relationships within the *Danio* genus showed markedly low support. The placement of *D. choprae* in the unpartitioned genomic topology was supported by 63% of jackknife windows, but a large minority, 27% of windows, placed *D. choprae* basal to the *Danio rerio* species group plus the *D. albolineatus*+*D. kerri* group (Fig. 2A). Analysis of chromosome structure showed that the placement of *D. choprae* tended to match the unpartitioned genomic topology at chromosome centers with *D. choprae* as sister to *D. erythromicron+D.margaritatus* but matched the alternative placement as sister to the *D.* rerio group + the *D. albolineatus+D. kerri* group at chromosome ends (Fig. 2D). A similar anomaly was observed for the placement of *D. nigrofasciatus*. The unpartitioned genome topology placed *D. nigrofasciatus* basal to *D. rerio, D. aesculapii,* and *D. kyathit.* This placement was supported by 71% of jackknife trees with a considerable bias for the center of chromosomes, while an alternative topology placed *D. nigrofasciatus* as sister to *D. kyathit* with support from 28% of windows occurring almost exclusively at chromosome ends (Fig. 2A, C; Fig. 3A).

**Figure 3:**
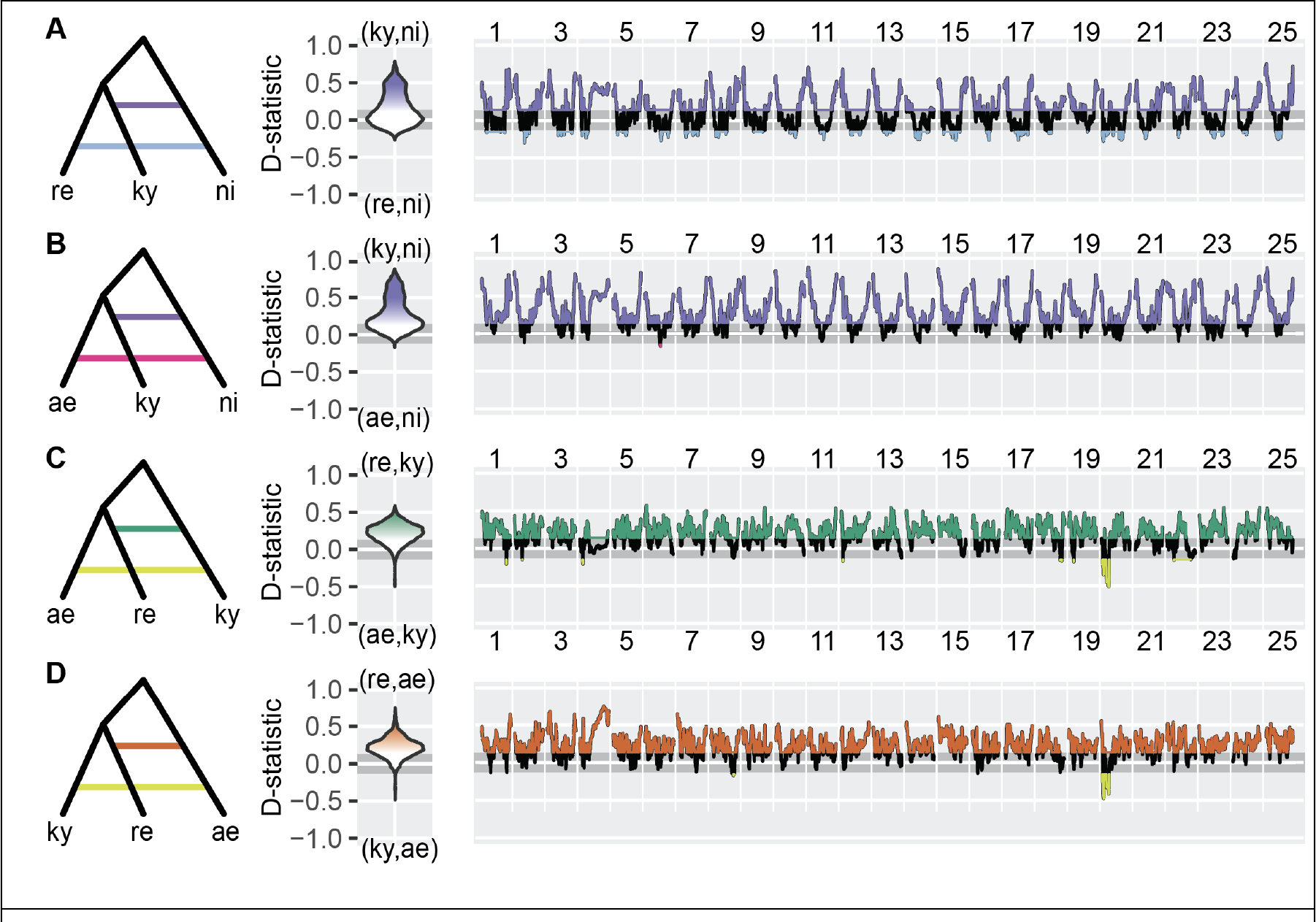
D-statistics show genomic regions affected by introgression. (A) D- statistics testing for gene flow between *D. nigrofasciatus* and *D. kyathit* (purple) or between *D. nigrofasciatus* and *D. rerio* (blue). The assumed topology (left), distribution of *D*-statistic values across the genome (middle column), and *D* values across chromosomes for windows of 200 ABBA/BABA sites (right, note the spike near telomeres). The horizontal gray bar shows the 95% confidence intervals under the null expectation of equal ABBA/BABA frequencies. Color scheme as in Fig. 2. (B) D- statistics testing for gene flow between *D. nigrofasciatus* and *D. kyathit* (purple) or between *D. nigrofasciatus* and *D. aesculapii* (pink). (C) D-statistics testing for gene flow between *D. kyathit* and *D. rerio* (green) or between *D. kyathit* and *D. aesculapii* (yellow), demonstrating an anomaly on Chr20, but no consistent effect of chromosome position relative to telomeres. (D) D-statistics testing for gene flow between D. aesculapii and *D. rerio* (rust) or between *D. aesculapii* and *D. kyathit* (yellow), again showing an anomaly on Chr20.

The placement of all species in the unpartitioned genomic topology was supported by the majority of the genome with one exception—the placement of *D. rerio*. Many (40%) of jackknife windows recovered *D. rerio* as sister to *D. aesculapii* (Fig. 2B) in agreement with the unpartitioned genomic topology, but a slight plurality of jackknife windows (42%) placed *D. rerio as* sister to *D. kyathit* (Fig. 2A). The few remaining trees placed *D. rerio* basal to two or more species in the *D. rerio* species group (Fig. 2A, C). The finding that *D. kyathit* and *D. aesculapii* have nearly equal support to be the sister of *D. rerio* is consistent with the origin of *D. rerio* by hybridization associated speciation.

The effect of genome structure on the history of the *D. rerio* genome is not subtle. *D. rerio* was recovered as sister to *D. kyathit* in 72% of the 50 jackknife partitions from the centers of chromosomes (Fig. 2C left, green), but in 0% of the 50 partitions from chromosome ends. In contrast, at chromosome ends, *D. rerio* was placed as sister to *D. aesculapii* in 58% of partitions (Fig. 2C right, rust) or diverging more basally in the *D. rerio* species group (42% of partitions, Fig. 2C right, purple). Notably, the relationships within the *D. rerio* species group recovered by the unpartitioned genomic topology match only 22% of jackknife windows (Fig. 2C), demonstrating that the unpartitioned genomic topology is insufficient to fully explain the history of the *D. rerio* species group.

### Genomic Regions Affected by Gene Flow and Effects of Recombination

To determine the sources of phylogenetic discordance within genus *Danio*, we focused on the most variable part of the phylogeny—the *D. rerio* species group—and patterns of pairwise splits, a subset of SDCs in which a derived character state is found in only two species. In a well-supported phylogeny with three species and an outgroup (Fig. 1), pairwise splits include BBAA splits shared by the most-closely related species, as well as ABBA and BABA splits (see Fig. 1). Several tests can use these “ABBA/BABA sites” to detect and quantify introgression and phylogenetic discordance (Martin, et al. 2015b; Pease and Hahn 2015; Blischak, et al. 2018; Kubatko and Chifman 2019).

To determine if introgression could explain the 28% of the jackknife trees that recovered *D. nigrofasciatus* as sister to *D. kyathit*, we calculated Patterson’s D-statistics genome-wide (Table 1) and in windows of 200 ABBA/BABA sites across the genome (Fig. 3). Genome-wide D-statistics revealed a striking excess of characters supporting introgression between *D. kyathit* and *D. nigrofasciatus* using either *D. rerio* (D = 0.14) or *D. aesculapii* (D = 0.29) as the sister species to *D. kyathit*. Although ABBA/BABA patterns were roughly equal (D not significantly different from 0) over much of the genome, the ends of chromosomes harbored an excess of derived characters shared by *D. kyathit* and *D. nigrofasciatus* (Fig. 3A, B; purple). This pattern of high introgression signal near the ends of chromosomes suggests introgression limited to the high recombination regions of the danio genome and explains why jackknife trees placing *D. kyathit* with *D. nigrofasciatus* were found almost exclusively at the ends of chromosomes (Fig. 2C).

**Table 1.** Genome-wide D-statistics within the *D. rerio* species group.

| Assumed Species Tree (P1,P2),P3 | *D*-<br>statistic | BBAA<br>sites | ABBA<br>sites | BABA<br>sites |
| --- | --- | --- | --- | --- |
| *(D.rerio,D.kyathit),D.nigrofasciatus* | 0.1405 | 63343 | 48670 | 36676 |
| *(D.aesculapii,D.kyathit),D.nigrofasciatus* | 0.2978 | 46262 | 49991 | 27046 |
| *(D.aesculapii,D.rerio),D.kyathit* | 0.2027 | 56970 | 54596 | 36194 |
| *(D.kyathit,D.rerio),D.aesculapii* | 0.223 | 54596 | 56970 | 36194 |

We similarly used D-statistics to test for evidence of admixture in *D. rerio*. Because two species—*D. aesculapii* and *D. kyathit—*had nearly equal support as the sister species of *D. rerio* (Fig. 2B), we calculated D-statistics for both possible relationships. Genome-wide D values were high for both comparisons (Table 1), providing strong support for an effect of admixture from both species on the history of the *D. rerio* genome. In stark contrast to the elevated D-statistics between *D. kyathit* and *D. nigrofasciatus*, which occurred almost exclusively at the ends of chromosomes (Fig. 3A,B), D-statistics grouping *D. rerio* with *D. kyathit* and *D. aesculapii* were elevated across the majority of the genome (Fig. 3C,D), supporting genome-wide admixture. These D-statistic tests, based on more than 80,000 ABBA/BABA sites per comparison, robustly reject ILS as the only source of variance in the *D. rerio* species group. Moreover, the genome-wide D values are greater than 0.14 in all comparisons, making them higher than in some other studies where introgression was either known *a priori*, or validated with follow up studies (Cui, et al. 2013; Sankararaman, et al. 2014; Li, et al. 2016). While D-statistic tests rule out ILS as the only source of genealogical discordance in danios, these tests do not distinguish between introgression and HAS.

To supplement results from Patterson’s D-Statistic tests and to compare all four species of the *D. rerio* group, we turned to a more generalized approach to investigate pairwise splits while also incorporating genomic position. Our dataset contains over 200,000 pairwise splits that inform relationships within the *D. rerio* species group, allowing for a more fine-scale investigation than the jackknife trees of Figure 2. If four species are all equally closely related, we expect to see each of the six pairwise splits occurring at equal frequency (16.7%). Regions of the genome having pairwise splits in considerable excess of this frequency support a closer relationship of the two species sharing the derived characters. Using a sliding window of 500 ancestry-informative SNPs, we find that each chromosome is a patchwork of genomic regions each enriched for a combination of pairwise splits (Fig. 4A, Supplemental Fig. S1).

**Figure 4:**
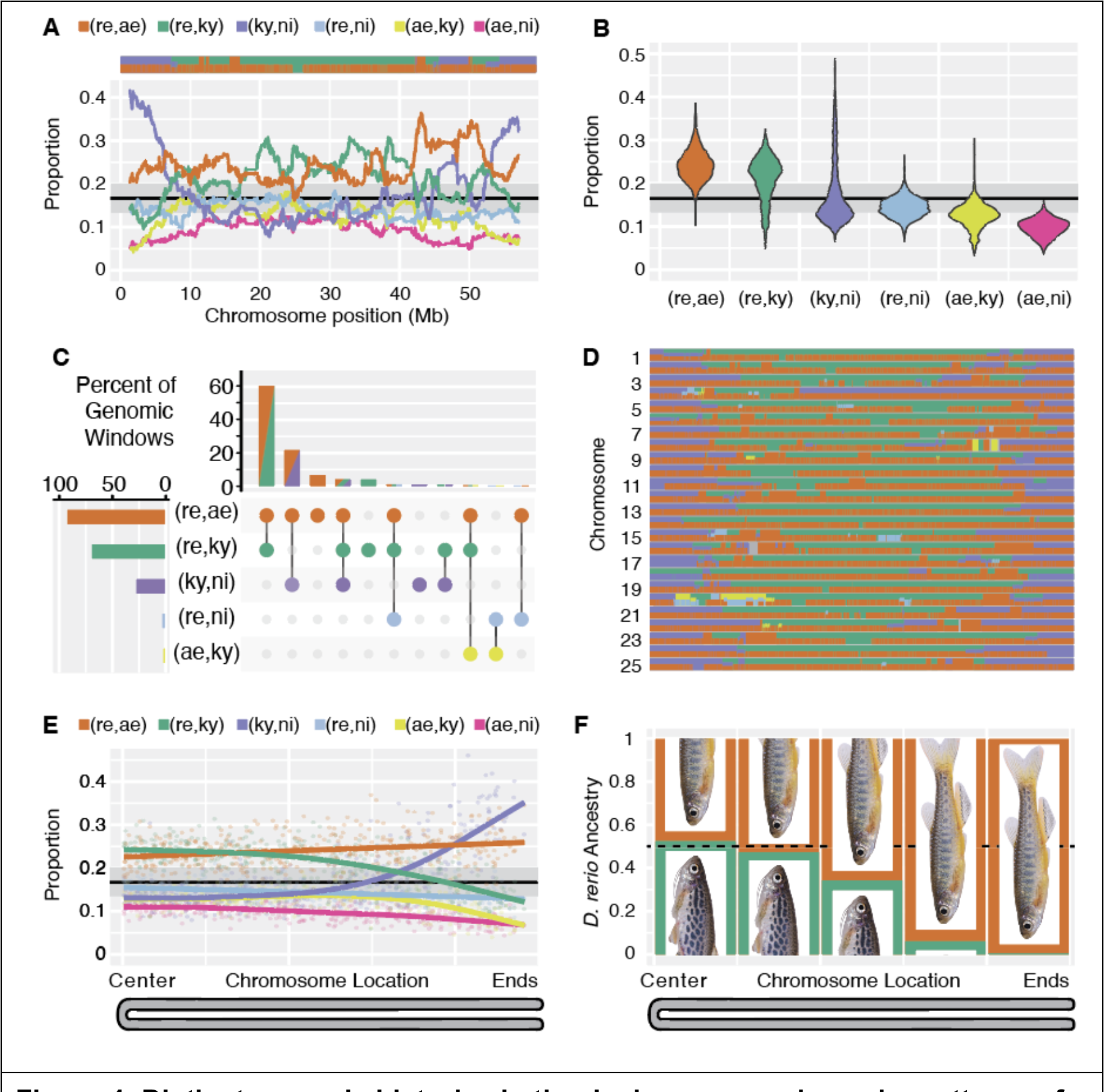
D**i**stinct **genomic histories in the danio genome shown by patterns of shared derived characters.** (A) Proportions of the six pairwise splits in the *D. rerio* species group in sliding windows of 500 pairwise splits across chromosome 2, which displays a representative pattern, and all chromosomes appear in Supplemental Fig. S2. The gray horizontal bar shows the 95% confidence interval under the null expectation of equal proportions of the six split patterns. The colored bar at the top shows which splits are enriched at each position along the chromosome. Color scheme as in Fig. 2. (B) Distribution of pairwise split proportions from the entire genome (windows of 500 pairwise splits). (C) Percentage of genomic windows showing enrichment for pairwise splits in at least 10 genomic windows. The bottom left graph shows the percent of the genome enriched for each combination of pairwise splits. (D) Co-occurrence of pairwise split enrichment across all 25 zebrafish chromosomes. Genomic regions with no significant enrichment are dark grey. (E) Effects of chromosome location on pairwise split proportions. Points are plotted according to relative chromosome location of a window along a folded chromosome (centers of chromosomes on the left and ends of chromosomes on the right) and the proportion of each pairwise split in that window. Splines are fitted to the data for each pairwise split across all 25 chromosomes in non-overlapping windows of 500 pairwise splits. (F) Proportion of *D. rerio* ancestry attributable to the *D. aesculapii* lineage (orange) or the *D. kyathit* lineage (green). Analysis was performed on the sequences used previously for jackknife trees binned according to chromosome position.

Genome-wide, all six pairwise splits appeared at appreciable frequency (Fig. 4B), suggesting a major role for ILS as danios speciated. Enrichment in genomic windows, however, was restricted almost exclusively to the three most common splits (re,ae; re,ky; ky,ni), which corresponded to relationships supported by both the jackknife trees (Fig. 2C) and the genome-wide *D*-statistics (Table 1).

The majority of genomic windows (65.2%) indicated that *D. rerio* is closely related to both *D. aesculapii* and *D. kyathit* (orange/green bars in Fig. 4C), suggesting that zebrafish arose from an ancient HAS event between ancestors of these two lineages. A further 10.4% of the genome indicated *D. rerio* was closely related to either *D. aesculapii* (6.4%) or *D. kyathit* (4.0%), but not the other species. A smaller proportion of windows (25.7%) indicated that *D. rerio* and *D. aesculapii* are closely related and that *D. kyathit* and *D. nigrofasciatus* are closely related (orange/purple bars in Fig. 4C). This set of relationships had been previously recovered with strong support (McCluskey and Postlethwait 2015). The separation of the danio genome into regions with two distinct histories is remarkably consistent when viewed across chromosomes, although small regions supporting other histories are apparent (Fig. 4D, Supplemental Fig. S2).

A small minority of genomic regions stood out in comparison to other chromosomes. One region spanning fifteen megabases on the left half of chromosome 20 supported a history unique from the rest of the genome (Fig. 4D; Supplemental Fig. S1). This region was enriched for pairwise splits grouping *D. rerio* with *D. nigrofasciatus* (blue in figures) and *D. aesculapii* with *D. kyathit* (yellow in figures). Several other instances of discrete regions with distinct histories have been described when chromosomal inversions have a paraphyletic distribution relative to the species tree (Hohenlohe, et al. 2010; Hohenlohe, et al. 2012; Fontaine, et al. 2015; Li, et al. 2016; Pease, et al. 2016).

Furthermore, studies of cyprinid karyotypes suggest that pericentromeric inversions are common in cyprinids (Avise and Gold 1977). The hypothesis that the patterns on Chromosome 20 correspond to one or more structural changes in the evolution of danios predicts that the gene order along the chromosome should differ between danio species.

The sex chromosome also showed a divergent pattern. SNPs grouping *D. kyathit* and *D. nigrofasciatus* were enriched at the ends of all 25 chromosomes (purple in Fig. 4D; Supplemental Fig. S2), but occurred at much lower frequency in the centers of all chromosomes except Chromosome 4, which contains a major sex determining gene in nature, with ZZ individuals always becoming male and ZW individuals usually developing as females, but with some sex reversal into neomales (Anderson, et al. 2012; Wilson, et al. 2014a; Valdivieso, et al. 2022; Wilson, et al. 2024; Wilson and Postlethwait 2024). The right arm of Chromosome 4 is non-recombining, repeat rich, and full of duplicated gene families, except for several megabases near the right telomere containing the sex locus, (Anderson, et al. 2012; Howe, et al. 2013; Wilson, et al. 2014b). Despite spanning tens of megabases, because it does not recombine, the right arm of Chromosome 4 is effectively a single locus near the end of the chromosome and thus is expected to match the history of chromosome ends, as the data showed (Fig. 4D).

Genome-wide, the two major histories of the *D. rerio* species group are partitioned according to chromosome location (Fig. 4D, E). The first major history was apparent in the centers of chromosomes and supported a hybrid origin of *D. rerio* as evidenced by an excess of splits shared by *D. rerio* and *D. kyathit* and an excess of splits shared by *D. rerio* and *D. aesculapii*. Both splits occurred at nearly double the rate of splits exclusive to *D. kyathit* and *D. aesculapii*, demonstrating that the excess was not due to ILS. The second major history was apparent at the ends of chromosomes with *D. rerio* sister to *D. aesculapii* and *D. kyathit* sister to *D. nigrofasciatus*. Notably, as splits grouping *D. kyathit* and *D. nigrofasciatus* became more common toward the ends of chromosomes, splits grouping *D. kyathit* and *D. rerio* became less common. This anticorrelation suggests that introgression between *D. kyathit* and *D. nigrofasciatus* supplanted many derived characters exclusive to *D. kyathit* and *D. rerio*, thereby obscuring the relationships between the latter two species.

Several analyses herein suggested that *D. rerio* represents a hybrid species between the *D. aesculapii* and *D. kyathit* lineages. To test this hypothesis, we used the program HyDe (Blischak, et al. 2018), which can detect evidence of hybridization and also estimate the fraction of ancestry attributable to each ancestral lineage. Using the 15.44 MB of aligned coding sequence used previously for jackknife analyses (Fig. 2C), the hybrid detection results were clear and striking (Fig. 4F; Supplemental Table S3). The central 40% of the zebrafish genome strongly supported a HAS origin of *D. rerio* (p < 10e-10), with 50% ancestry estimates each for *D. aesculapii* and *D. kyathit.* The apparent contribution of the *D. kyathit* lineage, however, declined towards the ends of chromosomes with <1% ancestry estimated from *D. kyathit*. This finding suggests that alleles contributed by the *D. kyathit* lineage to the ends of chromosomes are no longer present in modern *D. kyathit*, possibly due to the high levels of introgression with *D. nigrofasciatus* seen at chromosome ends (Fig 3A,B, Fig 4E).

## Discussion

### Admixture in the history of the Danio genus

This genome-wide analysis of exome sequences across ten species of the genus *Danio* provides a novel understanding of the origins of the zebrafish *Danio rerio*, a major biomedical model organism (Baldridge, et al. 2021). Results showed that the genomes of zebrafish and its three closest allies harbor two histories that have remarkably consistent distributions across all chromosomes and a third history restricted to a small portion of a single chromosome arm. The genomic distributions of these disparate histories point to major effects of introgression and hybrid speciation within this group (Fig. 5).

**Figure 5:**
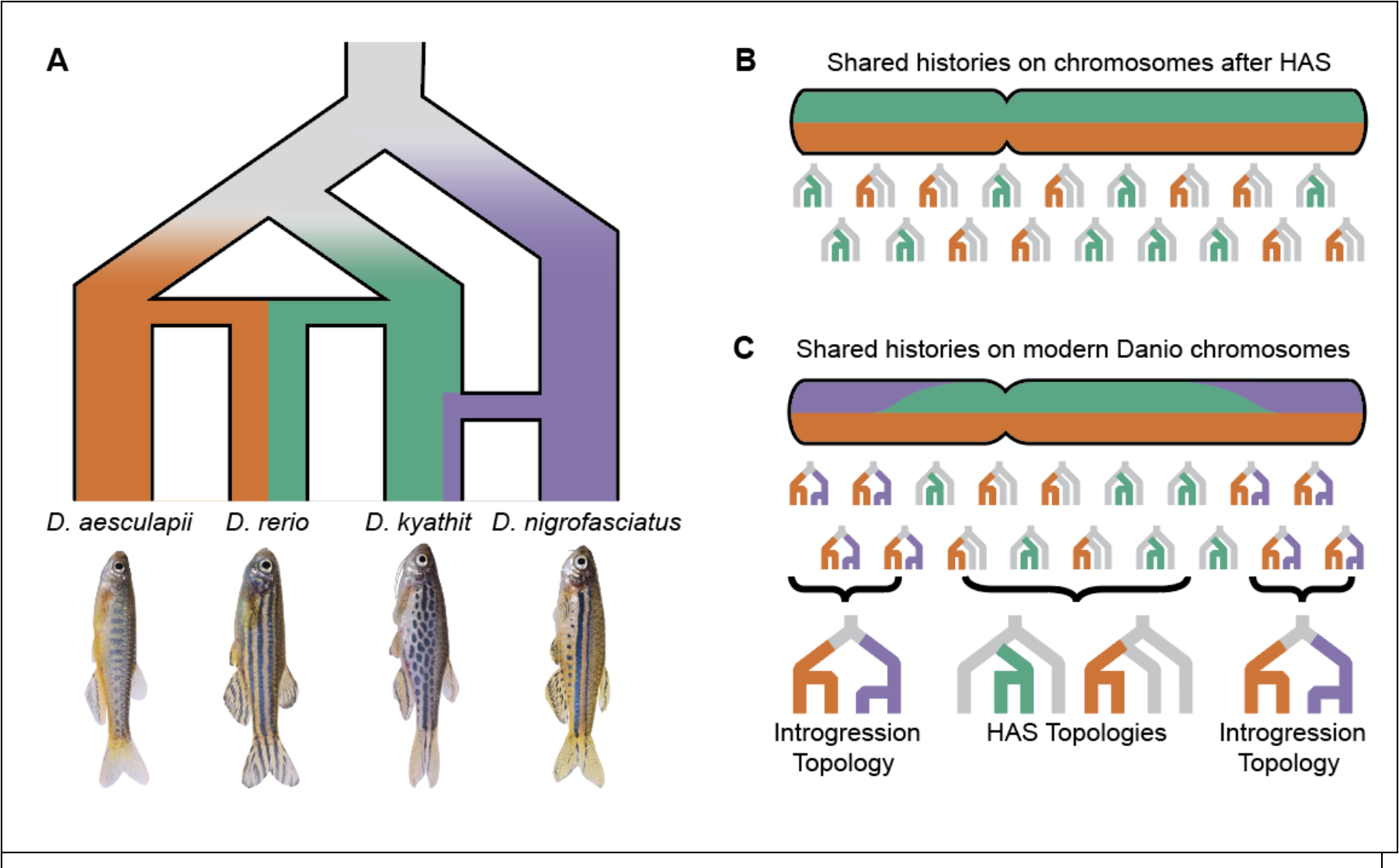
T**h**e **hybrid history of the zebrafish genome.** (A) A model for the recent population history of the *D. rerio* species group showing the hybrid origin of *D. rerio* from the *D. aesculapii* and *D. kyathit* lineages and the introgression of sequences between *D. kyathit* and *D. nigrofasciatus* retained mostly at chromosome ends. (B) Graphic depicting, across a chromosome, the expected distribution of alleles originating in populations ancestral to only two species at the time of hybridization associated speciation. (C) Approximate experimentally discovered distribution of alleles from populations shared by only two species in modern day danios.

The first history we observed is that *D. rerio* is closely -- and about equally -- related to both *D. aesculapii* and *D. kyathit* across the majority of the genome, while the latter two species are more distantly related to each other. This history is supported by phylogenies inferred from jackknife windows (Fig. 2), D-statistics, (Fig. 3), and the correlated genomic distributions of derived character states (Fig. 4C, E; Supplemental Fig. S1). All these results point to an ancient HAS origin of *D. rerio* from ancestors in the *D. aesculapii* and *D. kyathit* lineages (Fig. 5). The second history we observed occurred in the regions of high recombination at the ends of danio chromosomes (Fig. 2E, Fig. 5). These locations showed a striking excess of derived alleles shared by *D. kyathit* and *D. nigrofasciatus,* consistent with introgression limited to high recombination regions (Fig. 4D,E; Supplemental Fig. S1). The third history is limited to a portion of chromosome 20, where *D. rerio* is not closely related to either *D. aesculapii* or *D. kyathit*, although the latter two species are closely related to each other in this region (Fig. 3D; Supplemental Fig. 3). A distinct history restricted to a single genomic region is consistent with a chromosomal inversion either inherited by ILS or passed by introgression as has been seen in other taxa (della Torre, et al. 1997; Kulathinal, et al. 2009; Fontaine, et al. 2015; Barron, et al. 2019; Edelman, et al. 2019; Nelson, et al. 2021; Soudi, et al. 2023).

These distinct histories point to a hybrid origin of *D. rerio*, but is this hybrid beginning due to derivation by a homoploid hybridization event? Opinions differ as to the criteria that define a homoploid hybrid species (Schumer, et al. 2014; Nieto Feliner, et al. 2017), but three proposed requirements include showing that: 1) the species is reproductively isolated from parent species; 2) genetic evidence shows hybridization; and 3) hybridization was responsible for the reproductive isolation (Schumer, et al. 2014).

Several studies have reported hybrid sterility in hybrids between *D. rerio* and other danios (Wong, et al. 2011;Podobnik, et al. 2020;Parichy and Johnson 2001), likely caused by aneuploid gametes (Endoh, et al. 2020). Analysis of exon sequences discussed here provide ample evidence that the *D. rerio* genome shows evidence of hybridization between the *D. aesculapii* and *D. kyathit* lineages. Whether hybridization was responsible for the reproductive isolation of *D. rerio* remains uncertain, however, and may be difficult to determine given one of the putative parental lineages shows evidence of subsequent gene flow. *D. rerio* may be a homoploid hybrid species, but further studies will be needed to understand better the origin of this model species.

Danios, and perhaps Cyprinids at large, offer a unique window into recombination variation and genomic histories because the group is remarkably diverse and clear patterns of recombination variation are present within each chromosome. The effects of recombination rate have been noted in several species across inversions, in sex chromosomes, and near recombination hot spots (Turner, et al. 2005; Noor and Bennett 2009; Turner and Hahn 2010). Theory predicts that regions of lower recombination rates, such as the centers of danio chromosomes, will be resistant to gene flow (Felsenstein 1974; Hill and Robertson 2007). This distribution would occur especially in the case of pericentric inversions, which cytogenetic studies suggest are common among cyprinids (Avise and Gold 1977). Conversely, regions with high recombination rates, such as the ends of danio chromosomes, will be more susceptible to gene flow following the beginning of reproductive isolation and should better reflect the phylogeographic distribution of species. Interestingly, relationships at the ends of danio chromosomes more closely reflect present geographical distributions of danios than do relationships at the centers of chromosomes. This pattern holds for *D. kyathit* and *D. nigrofasciatus*, which both occur in the Irrawaddy hydrological basin; for *D. rerio* and *D. aesculapii*, which both occur in Bangladesh; and for *D. choprae*, which does not occur with *D. erythromicron* or *D. margaritatus* but does occur in the same drainage as several other danio species (McCluskey and Postlethwait 2015).

Much remains to be learned about the history of danios and the hybrid history of zebrafish. Inferences drawn here were based on ten *Danio* species, but the study of more danio species will help us better understand these relationships. Notably, two members of the *D. rerio* species group -- the striped *D. quagga* and the spotted *D. tinwini* -- may prove to be the best representatives of the lineages that experienced introgression in this group (Kullander and Fang 2009b; Kullander, et al. 2009). Finally, the mitochondrial phylogeny for these species could not be inferred in this study because the exome enrichment baits targeted only nuclear genes. A better understanding of mitochondrial histories will provide insight into the history of these species.

With current taxon sampling and topologies of the best supported trees, we cannot determine the direction of introgression in these species using conventional methods, which require comparison of two pairs of sister species (Durand, et al. 2011; Martin, et al. 2015a; Pease, et al. 2016) or estimates of historical effective population size (Hibbins and Hahn 2022). Future analyses incorporating more taxa, population level sampling across species, and whole mitochondrial and nuclear genome analyses may help to resolve better the complex history of members of the *Danio* genus.

### Implications for Future Studies

The structured distribution of relationships across the danio genome has implications for the interpretation of past and future phylogenetic studies. First, when different genomic regions have different histories, the use of a small number of loci for phylogenetic inference will have considerable limitations. Previous phylogenetic studies involving the *Danio* genus (Mayden, et al. 2007; Fang, et al. 2009; Tang, et al. 2010; McCluskey and Postlethwait 2015) arrived at different conclusions, likely in part because the loci used for inference were different in different studies and were sometimes sampled from chromosome regions that our data show to have different histories. The two nuclear markers most frequently used in these species are *rhodopsin* near the end of chromosome 8, and *rag1* closer to the center of chromosome 25, regions we show tend to have different histories. Second, when large phylogenomic datasets are used without incorporating data on genome structure, systematic biases can lead to different inferred relationships than when genome structure is incorporated into analyses. For example, our previous phylogenomic study of *Danio* used short sequences flanking *SbfI* restriction sites sampled across the genome (McCluskey and Postlethwait 2015). Using a variety of datasets and phylogenetic inference methods, the best-supported topology for the *D. rerio* group in that study placed *D. kyathit* with *D. nigrofasciatus* and *D. rerio* with *D. aesculapii*, a topology supported by only 16.8% of jackknife windows in the current exome study with an extreme bias toward the ends of chromosomes. An analysis of *SbfI* sites in the *D. rerio* genome showed that these restriction sites have an extreme bias for the ends of chromosomes (Supplemental Figure S2), which we show here to be greatly enriched for SNPs supporting the topology recovered in the previous study.

Interpretation of relationships in the context of genome structure will have important ramifications in future studies because drawing inferences under the supposition of different species trees can have considerable implications on inferred evolutionary origins of species-specific biology in the taxa examined (Thomas and Hahn 2015). Indeed, without accounting for genome structure, the history of the *D. rerio* species group inferred from the entire genome (the unpartitioned genomic topology) is supported by only 22.0% of genomic regions (Fig. 2C) and does not reflect the high levels of introgression between *D. kyathit* and *D. nigrofasciatus* or the hybrid origin of zebrafish.

## Methods

### Library preparation and Exome Alignment Creation

DNAs were collected from the following species: zebra danio (*Danio rerio*, AB strain), orange-finned danio (*Danio kyathit*), panther danio (*Danio aesculapii*), spotted danio (*Danio nigrofasciatus*), pearl danio (*Danio albolineatus*), Kerr’s danio (*Danio kerri*), glowlight danio (*Danio choprae*), celestial pearl danio (*Danio margaritatus*), emerald dwarf danio (*Danio erythromicron*), Meghalaya danio (*Danio meghalayensis*), and giant danio (*Devario aequipinnatus*). Fish were sourced from local hobby aquarium suppliers with the exception of *Danio rerio*, which were AB strain and sourced from the Zebrafish International Research Center (Eugene, OR). The University of Oregon Animal Care and Use Committee approved all protocols associated with this work. We extracted and purified genomic DNA using a Blood and Tissue Kit (Qiagen) and constructed libraries and performed exome enrichment with SureSelectXT2 RNA oligonucleotide baits (Agilent) according to manufacturer’s instructions. Exome-enriched libraries were quantified using a Qubit fluorimeter and sequenced on the Illumina HiSeq 2500 (Single End100 bp reads) and the Illumina NextSeq (Single End 75 bp reads). We quality-filtered Illumina reads with Trimmomatic (Bolger, et al. 2014) using a per-base quality score minimum set to 20 and minimum read length of 30 nucleotides.

To generate orthologous sequence alignments for phylogenetic inference, we first aligned genomic reads to coding sequence for APPRIS primary gene models (Rodriguez, et al. 2013) from the zebrafish genome GRCz10 v82 (Kersey, et al. 2016). We used GSNAP release 2015-07-23 (Wu and Nacu 2010) with the following parameters to account for mapping genomic reads to a transcript reference and for the sequence divergence between species: -k 12 --expand-offsets=1 --max-mismatches=0.2 --npaths=1. Consensus sequences for each species were generated with samtools v 1.1 (Danecek, et al. 2021) using the mpileup command with parameters: -B -C 0 -Q 0 -q 0 - m 1 -e 0 -F 0 -h 0 -o 0 followed by the “bcftools call” command with the -c flag. These parameters ensured that reads with any evidence of indels were flagged and excluded, while allowing soft clipping of the genomic reads to the reference transcripts. Consensus sequences from 11,635 genes each having at least 500 unambiguously aligned nucleotide positions with at least 10x coverage across each species were concatenated to form the 15.45 megabase unpartitioned, genome-wide data set. These aligned positions were split according to chromosome and position to create the 250 jackknife alignments.

To identify variants in danio species across the danio genome, we aligned Illumina reads for all 11 species to the zebrafish genome (GRCz10 v 82). To handle the amount of sequence divergence between the zebrafish reference genome and sequences from other species, we used bbmap (Bushnell 2014), a global alignment algorithm with permissive parameters, but required a single best alignment to the zebrafish genome. The BBMap parameters were: ambiguous=best minidentity=0.70 maxindel=100 idtag=t k=12. To extract variants from coding sequence across the genome, we used samtools mpileup with the following parameters: --no-BAQ --adjust-MQ 0 --min-BQ 13 --min-MQ 0 --min-ireads 1 --ext-prob 20 --gap-frac 0.002 --tandem-qual 100 --VCF --uncompressed - -output-tags DP,DPR,DV,DP4,INFO/DPR,SP. We then filtered variants using the “vcf-annotate” function from vcftools v0.1.12a to include only sites with at least 5x coverage and exclude sites with three or more SNPs within ten base pairs. The resulting variant calls were kept for subsequent analysis outlined below.

### Comparisons to Existing Sequences

Reconstructed sequences for *rho* and *rag1* from each species included in this study were compared to all available *Danio* (taxid:7954) and *Devario* (taxid:439832) sequences using BLASTN with default parameters on the NCBI BLAST site. The top BLAST hit to the appropriate species is included in Supplemental Table S2. In one instance (*D. meghalayensis rag1*), no sequences were available, so a match from the closely related *D. dangila* was included. When multiple matches were present, the match with the highest percent identity was selected.

### Maximum Likelihood Phylogenetic Analyses and Hybridization Detection

To infer phylogenies for different jackknife windows, we partitioned data from each chromosome into ten bins each representing ten percent of the total aligned sequence for each chromosome. For the unpartitioned genomic alignment and each of the 250 jackknife partitions, we inferred maximum likelihood phylogenies and approximate likelihood ratio tests under a GTR+I+**Γ** model in *RAxML* v 8.2.3 (Stamatakis 2006) on Talapas, the University of Oregon’s super computer (https://racs.uoregon.edu/talapas). We determined topology frequencies and visualized phylogenies using DensiTree (Bouckaert 2010). We performed concordance analyses in *BUCKy* v 1.4.2 (Larget, et al. 2010) with default settings.

To test our proposed model of a hybrid origin of *D. rerio* between the *D. aesculapii* and *D. kyathit* lineages, we used HyDe for hybrid detection (Blischak, et al. 2018). Sequence alignments analyzed were the same as those used for the 250 jackknife trees, partitioned according to chromosome position. Taxa were input such that *D. nigrofasciatus* was the outgroup and the value of gamma returned by HyDe corresponded to the proportion of the *D. rerio* genome with ancestry coming from the *D. kyathit* lineage.

### Genomic Structure analyses

To investigate relationships along the genome at a finer scale, we extracted genotypes from .vcf files for the genomic alignments outlined previously. We recoded genotypes (A,C,G or T) at biallelic sites to splits (A, ancestral, or B, derived) for subsequent analysis and visualization. To avoid confounding factors of regions of exceptionally low diversity, split frequencies were calculated for variable positions only. Due to variation in gene density and the discontiguous nature of our exome data across the genome, we standardized analyses to include the same number of splits rather than analyzing relative to windows of equal numbers of nucleotides in the nuclear genome. This approach ameliorated the effects of stochastic sampling that can cause false positives in certain types of genome scans (Martin, et al. 2015a). Chromoplots and line plots were plotted in R using the plot function of the ggplot2 package.

D-statistics were calculated as previously described (Durand, et al. 2011) using windows of 200 ABBA/BABA sites and plotted for every site based on the calculated value from the surrounding 200 ABBA/BABA sites. Ancestral alleles were determined based on sequences for all available basal danios. Any sites with polymorphism between basal species were removed to avoid the possible influence of introgression with the outgroup. For the analysis of all six pairwise splits, sliding windows of 500 splits, jumping by 10 sites were used. Regions enriched for particular splits were identified in R using a binomial distribution with an expected frequency of one sixth (based on six possible species pairs in the *rerio* species group) and the number of trials set to the window size of 500. RAD-site locations were from (McCluskey and Postlethwait 2015). Gene locations were downloaded from Ensembl BioMart for protein-coding genes on the 25 chromosomes from GRCz10 v 82.

---

**Supplemental Table S1:** Experiment Summary.

| Species | Total Reads | Reads |  |  |
| --- | --- | --- | --- | --- |
|  |  | Aligned to | Proportion | Exome |
|  |  | GRCz10 | Aligned | Coverage |
| *Danio rerio* | 41603616 | 41292106 | 0.993 | 87.839 |
| *Danio aesculapii* | 35171505 | 31812385 | 0.904 | 67.673 |
| *Danio kyathit* | 50412319 | 45793130 | 0.908 | 97.414 |
| *Danio nigrofasciatus* | 37389413 | 33110288 | 0.886 | 70.434 |
| *Danio albolineatus* | 31916307 | 27771158 | 0.870 | 59.077 |
| *Danio kerri* | 39820645 | 34829199 | 0.875 | 74.091 |
| *Danio choprae* | 52038977 | 45517922 | 0.875 | 96.829 |
| *Danio erythromicron* | 25080033 | 20885438 | 0.833 | 44.429 |
| *Danio margaritatus* | 35758836 | 30482592 | 0.852 | 64.845 |
| *Danio meghalayensis* | 39328467 | 33221487 | 0.845 | 70.671 |
| *Devario aequipinnatus* | 31491855 | 22986682 | 0.730 | 48.899 |

**Supplemental Table S2:** BLAST results for reconstructed *rho* and *rag1* sequences compared to publically available *Danio* and *Devario* nucleotide sequences.

| Reconstructed Exome Sequence | Accession<br>number | Percent<br>Identity | Alignment<br>Length | Accession<br>Species | Description |
| --- | --- | --- | --- | --- | --- |
| Drer_rho_ENSDART00000027000 | LR812070.1 | 100 | 467 | Danio rerio | Danio rerio genome assembly, chromosome: 8 |
| Daes_rho_ENSDART00000027000 | EU241365.1 | 99.399 | 333 | Danio aesculapii | Danio sp. 'snakeskin' rhodopsin (Rho) gene, partial cds |
| Dkya_rho_ENSDART00000027000 | LR812534.1 | 99.786 | 467 | Danio kyathit | Danio kyathit genome assembly, chromosome: 16 |
| Dnig_rho_ENSDART00000027000 | HM223909.1 | 100 | 317 | Danio nigrofasciatus | Danio nigrofasciatus isolate CTOL01597 rhodopsin gene, partial cds |
| Dalb_rho_ENSDART00000027000 | HQ286331.1 | 98.738 | 317 | Danio<br>albolineatus | Danio albolineatus voucher<br>7205 rhodopsin<br>photopigment (rh1) mRNA,<br>partial cds |
| Dker_rho_ENSDART00000027000 | JQ614125.1 | 100 | 320 | Danio kerri | Danio kerri rhodopsin (Rho)<br>gene, partial cds |
| Dcho_rho_ENSDART00000027000 | JQ614130.1 | 99.688 | 320 | Danio choprai | Danio choprai voucher<br>RC0060 rhodopsin (Rho)<br>gene, partial cds |
| Dery_rho_ENSDART00000027000 | HM223905.1 | 100 | 285 | Danio<br>erythromicron | Microrasbora<br>erythromicron isolate<br>CTOL00425 rhodopsin<br>gene, partial cds |
| Dmar_rho_ENSDART00000027000 | JQ614141.1 | 99.688 | 320 | Danio<br>margaritatus | Danio margaritatus<br>voucher RC0107 rhodopsin<br>(Rho) gene, partial cds |
| Dmeg_rho_ENSDART00000027000 | JQ614143.1 | 100 | 320 | Danio<br>meghalayensis | Danio meghalayensis<br>voucher RC0565 rhodopsin<br>(Rho) gene, partial cds |
| DEVaeq_rho_ENSDART00000027000 | JQ614155.1 | 98.125 | 320 | Devario<br>aequipinnatus | Devario aequipinnatus<br>voucher RC0352 rhodopsin<br>(Rho) gene, partial cds |
| ABZ_rag1_ENSDART00000142917 | LR812087.1 | 96.122 | 1083 | Danio rerio | Danio rerio genome<br>assembly, chromosome: 25 |
| Daes_rag1_ENSDART00000142917 | LR812502.1 | 96.399 | 1083 | Danio aesculapii | Danio aesculapii genome<br>assembly, chromosome: 25 |
| Dkya_rag1_ENSDART00000142917 | LR812543.1 | 96.214 | 1083 | Danio kyathit | Danio kyathit genome<br>assembly, chromosome: 25 |
| Dnig_rag1_ENSDART00000142917 | EU292699.1 | 96.781 | 994 | Danio<br>nigrofasciatus | Danio nigrofasciatus isolate<br>CTOL01597 recombination<br>activating protein 1 (RAG1)<br>gene, exon 3 and partial<br>cds |
| Dalb_rag1_ENSDART00000142917 | EU292696.1 | 96.329 | 1008 | Danio<br>albolineatus | Danio albolineatus isolate<br>CTOL01500 recombination<br>activating protein 1 (RAG1)<br>gene, exon 3 and partial<br>cds |
| Dker_rag1_ENSDART00000142917 | HM224029.1 | 96.925 | 1008 | Danio kerri | Danio kerri isolate<br>CTOL02792 recombination<br>activating protein 1 (RAG1)<br>gene, partial cds |
| Dcho_rag1_ENSDART00000142917 | HM224026.1 | 97.805 | 410 | Danio choprai | Danio choprai isolate<br>CTOL01667 recombination<br>activating protein 1 (RAG1)<br>gene, partial cds |
| Dery_rag1_ENSDART00000142917 | EU292698.1 | 98.28 | 407 | Danio<br>erythromicron | Microrasbora<br>erythromicron isolate<br>CTOL01953 recombination<br>activating protein 1 (RAG1)<br>gene, exon 3 and partial<br>cds |
| Dmar_rag1_ENSDART00000142917 | EU292695.1 | 98.034 | 407 | Danio<br>margaritatus | Danio margaritatus isolate<br>CTOL02780 recombination<br>activating protein 1 (RAG1)<br>gene, exon 3 and partial<br>cds |
| Dmeg_rag1_ENSDART00000142917 | EU292697.1 | 98.025 | 405 | Danio dangila | Danio dangila isolate<br>CTOL01501 recombination<br>activating protein 1 (RAG1)<br>gene, exon 3 and partial<br>cds |
| DEVaeq_rag1_ENSDART00000142917 | KJ548110.1 | 94.729 | 645 | Devario<br>aequipinnatus | Devario aequipinnatus<br>isolate 5 recombination<br>activating protein 1 (RAG1)<br>gene, partial cds |

**Supplemental Table S3:**
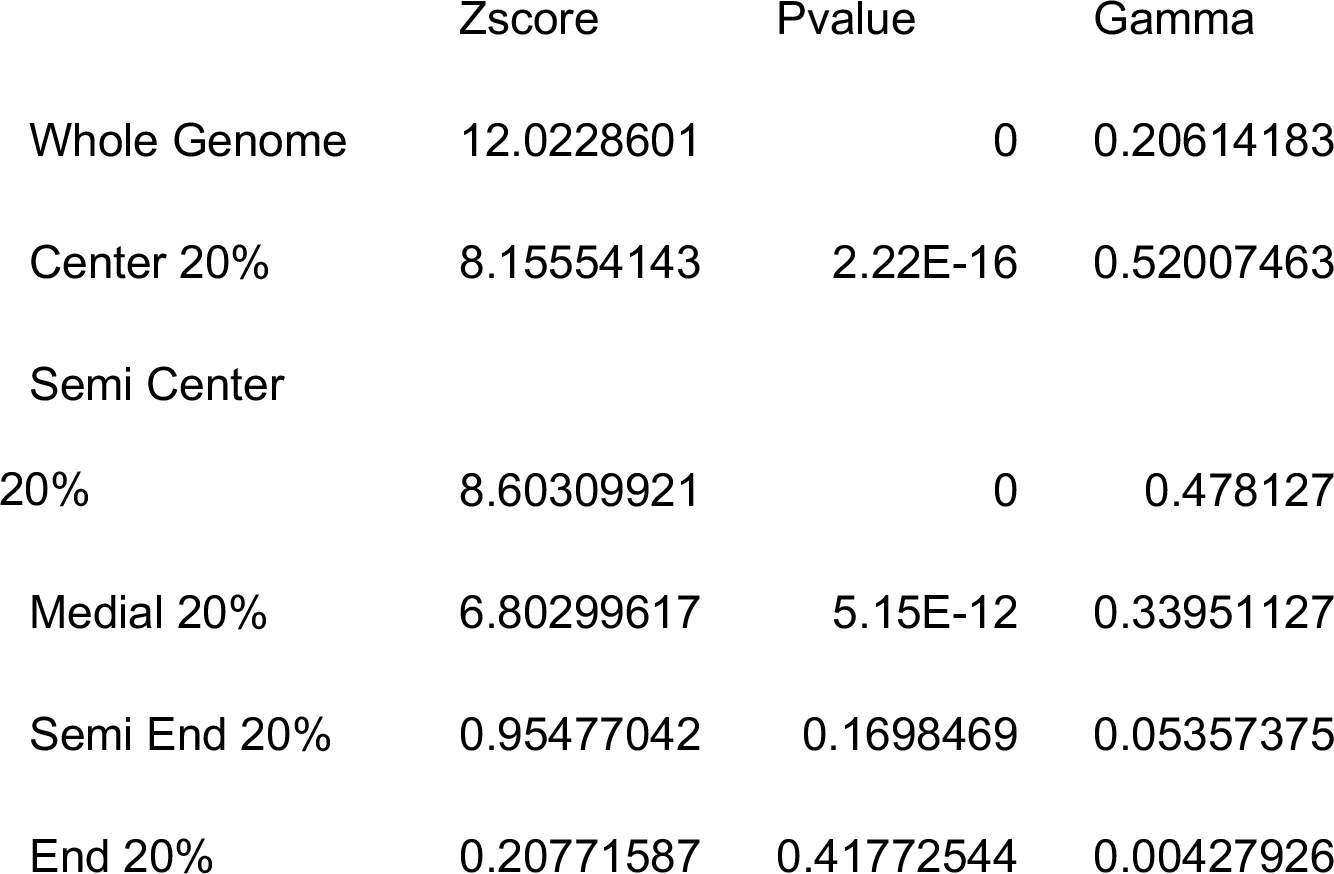
HyDe Analyses.

**Supplemental Figure S1:**
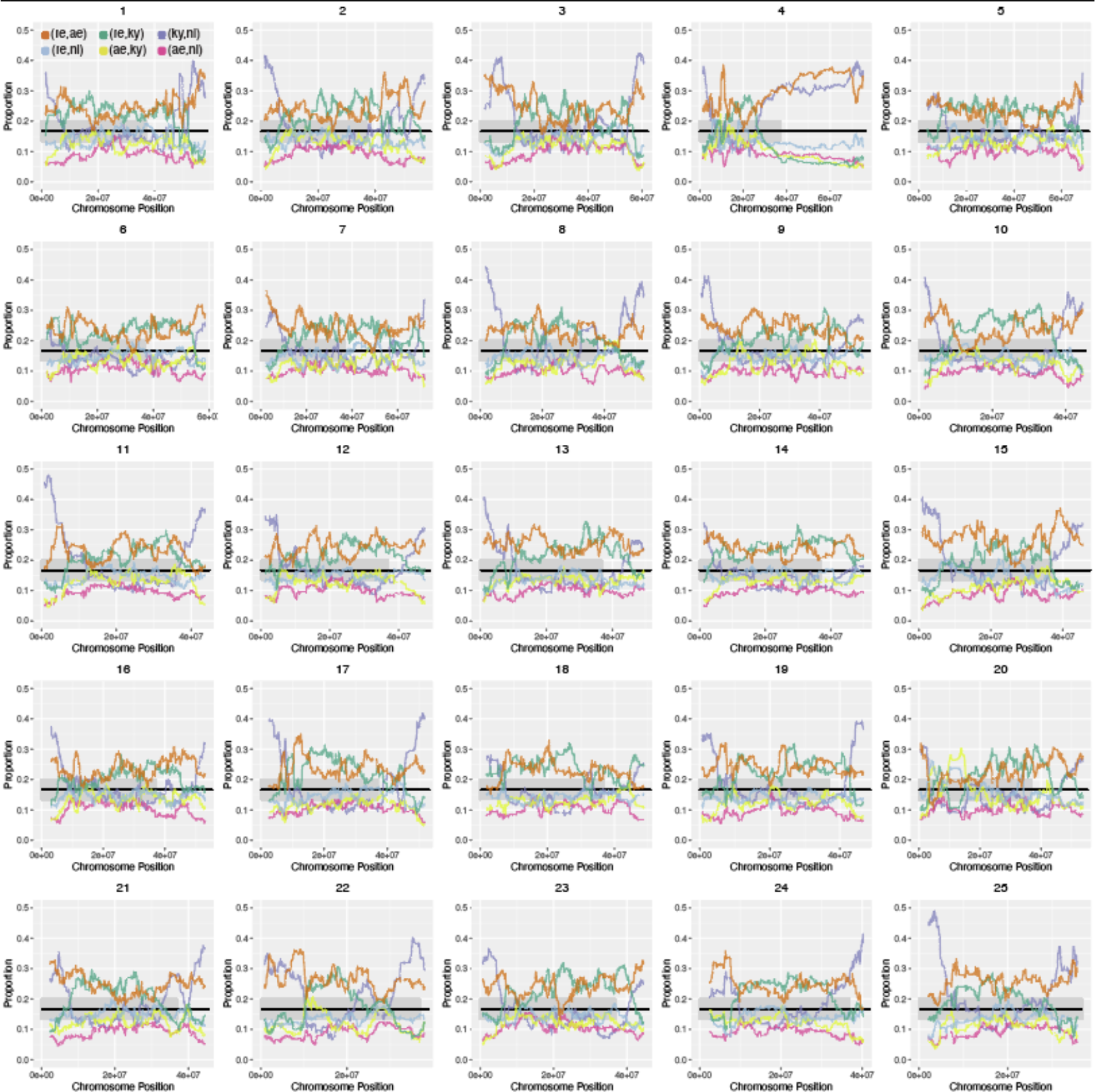
Pairwise split frequencies across each chromosome. Proportions of the six pairwise splits in the *D. rerio* species group in sliding windows of 500 pairwise splits across each chromosome. The gray bar shows the 95% confidence interval under the null expectation of equal split proportions.

**Supplemental Figure S2:**
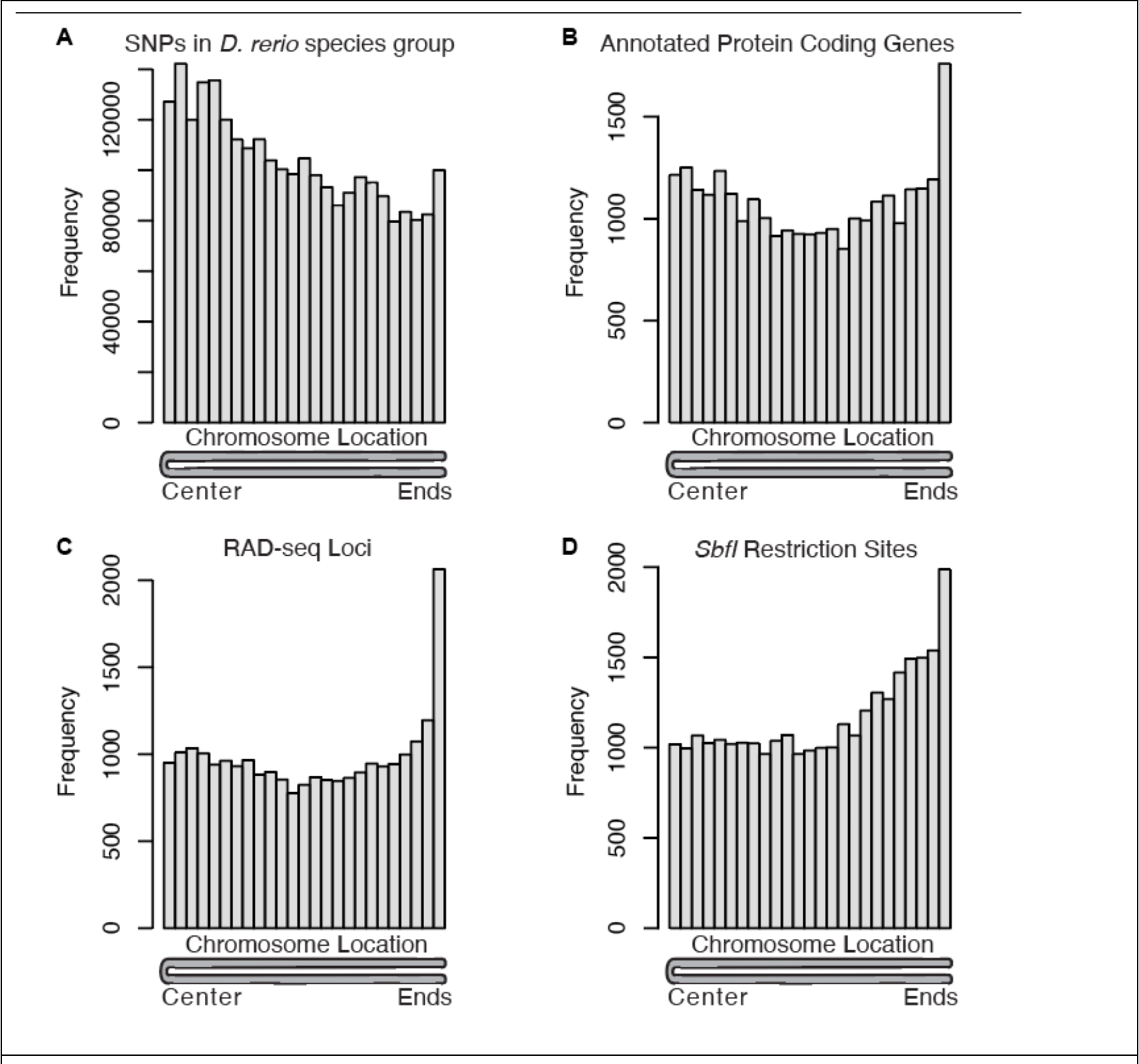
Genome-wide distribution of genomic features and data used for phylogenetic inference. (A) Histogram of SNPs in the *D. rerio* species group used in this study. Bins on the left are near the centers of chromosomes. Bins on the right are near the ends of chromosomes. (B) Histogram of annotated protein- coding genes in the zebrafish genome (GRCz10). (C) Histogram of RAD-seq loci used in (McCluskey and Postlethwait 2015). (D) Location of *SbfI* restriction sites in the zebrafish genome.

